# SARS-CoV-2 infection induces inflammatory bone loss in golden Syrian hamsters

**DOI:** 10.1101/2021.10.08.463665

**Authors:** Wei Qiao, Hui En Lau, Huizhi Xie, Vincent Kwok-Man Poon, Chris Chung-Sing Chan, Hin Chu, Shuofeng Yuan, Terrence Tsz-Tai Yuen, Kenn Ka-Heng Chik, Jessica Oi-Ling Tsang, Chris Chun-Yiu Chan, Jian-Piao Cai, Cuiting Luo, Kwok-Yong Yuen, Kenneth Man-Chee Cheung, Jasper Fuk-Woo Chan, Kelvin Wai-Kwok Yeung

**Author notes:** **Corresponding authors:** (J.F.-W.C.) and (K.W.-K.Y.).

## Abstract

Extrapulmonary complications of different organ systems have been increasingly recognized in patients with severe or chronic Coronavirus Disease 2019 (COVID-19). However, limited information on the skeletal complications of COVID-19 is known, even though inflammatory diseases of the respiratory tract have been known to perturb bone metabolism and cause pathological bone loss. In this study, we characterized the effects of severe acute respiratory syndrome coronavirus 2 (SARS-CoV-2) infection on bone metabolism in an established golden Syrian hamster model for COVID-19. SARS-CoV-2 causes significant multifocal loss of bone trabeculae in the long bones and lumbar vertebrae of all infected hamsters. The bone loss progressively worsens from the acute phase to the post-recovery phase. Mechanistically, the bone loss was associated with SARS-CoV-2-induced cytokine dysregulation which upregulates osteoclastic differentiation of monocyte-macrophage lineage. The pro-inflammatory cytokines further trigger a second wave of cytokine storm in the skeletal tissues to augment their pro-osteoclastogenesis effect. Our findings in this established hamster model suggest that pathological bone loss may be a neglected complication which warrants more extensive investigations during the long-term follow-up of COVID-19 patients. The benefits of potential prophylactic and therapeutic interventions against pathological bone loss should be further evaluated.

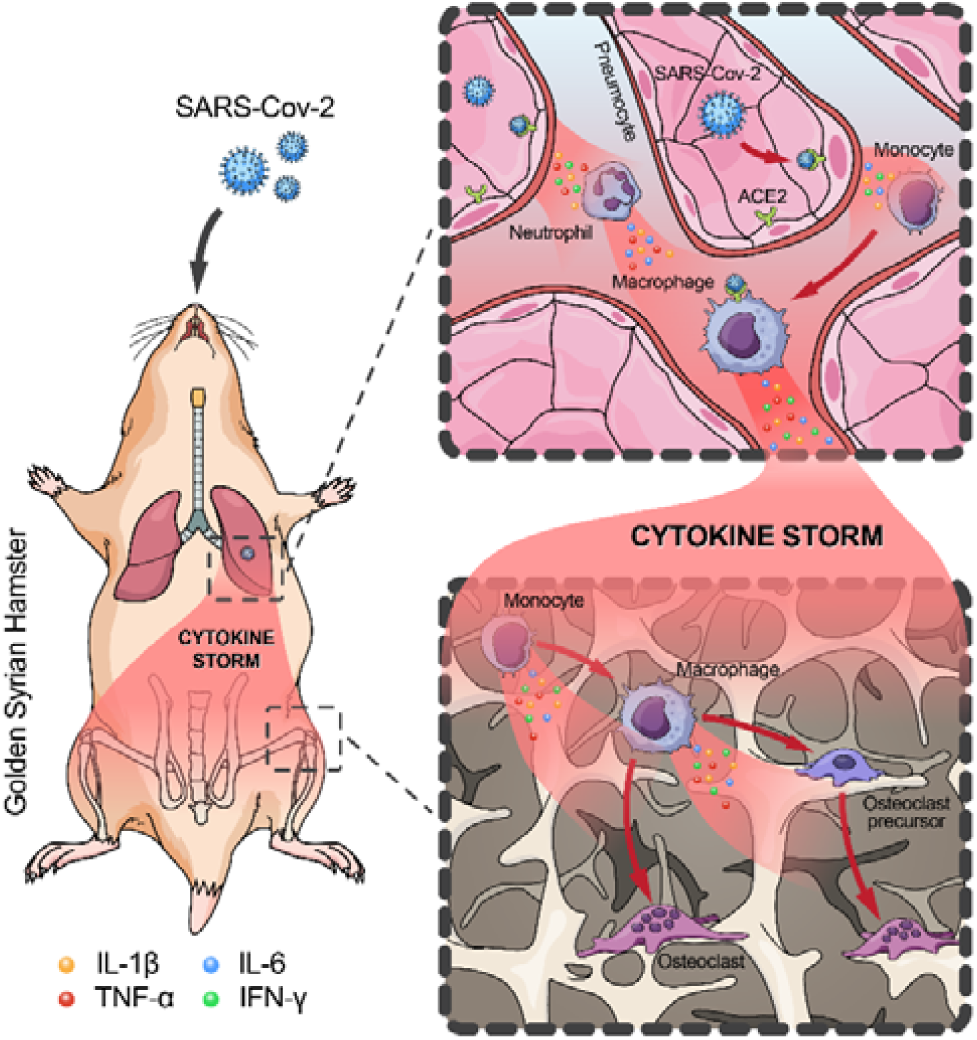

*Graphical abstract:* SARS-CoV-2 infection causes pathological bone loss in golden Syrian hamsters through induction of cytokine storm and inflammation-induced osteoclastogenesis.

## Introduction

The severe acute respiratory syndrome coronavirus 2 (SARS-CoV-2) has caused nearly 216 million cases of Coronavirus Disease 2019 (COVID-19) and nearly 4.5 million deaths as of 29 August 2021 since the virus’ discovery in December 2019 ^1^. Severe acute COVID-19 may be complicated by both pulmonary (pneumonia with acute respiratory distress syndrome and respiratory failure) and extrapulmonary manifestations, such as anosmia, ageusia, diarrhea, lymphopenia, and multi-organ dysfunction syndrome ^2–4^. More recently, it has been increasingly recognized that some patients may develop long-term complications and persistent symptoms of COVID-19, such as fatigue, headache, dyspnea, anosmia, muscle weakness, low-grade fever, and cognitive dysfunction ^5–7^. However, the full spectrum of clinical manifestations in the long-term post-acute sequelae of SARS-CoV-2 infection, or “long COVID”, remains incompletely understood. In particular, SARS-CoV-2-associated pathological changes of the skeletal system remain largely unknown.

Recently, a multi-centre study showed that COVID-19 patients requiring intensive care had significantly lower bone mineral density (BMD) than those who were managed in the non-intensive care setting ^8^. Another clinical study found that the number of severe clinical incidence was significantly higher in patients with lower BMD compared to those with higher BMD, therefore vertebral BMD is a strong independent predictor of mortality in COVID-19 patients ^9^. In addition, about 24% of long COVID patients reported bone ache or burning, with the symptoms lasting for up to 7 months after the onset of COVID-19 ^10^. Despite these emerging evidence on long-term complications of COVID-19, very limited serial investigations have been conducted on skeletal system involvement in the post-recovery phase. This is not unexpected because in COVID-19, patients either succumb or recover from the acute phase. In patients who recover from COVID-19, the focus on follow-up is usually limited to the respiratory, cardiac, and neurological functions which are well reported in the literature, rather than skeletal pathologies which typically do not manifest in the acute phase and therefore may be neglected. Moreover, since severe COVID-19 is most often found in elderly patients and those with comorbidities including patients those on chronic corticosteroid and immunosuppressive drugs, the virus-induced bone changes in these patients who may already have osteoporosis before the infection may not be appreciated.

The skeletal system undergoes continuous bone formation and degradation throughout life and this tightly regulated remodelling process could be disturbed by many factors, particularly metabolic alterations and hormonal changes ^11,12^. An imbalance between bone formation and resorption can also result from various chronic inflammatory diseases, leading to systemic osteoporosis and increased fracture risk ^13,14^. For instance, chronic pulmonary inflammation arising from chronic obstructive pulmonary disease (COPD), cystic fibrosis, and asthma are reported to induce systemic bone loss ^15,16^. Indeed, it has been shown that the extent of local or systemic osteopenia is associated with the degree of inflammatory response and the inflammation-induced bone loss can continue after effective therapeutic intervention on the inflammatory disease ^13,17,18^. Severe COVID-19 patients developed much higher serum concentrations of pro-inflammatory cytokines and chemokines (e.g., IL-1β, IFN-α, IL-1RA and IL-8) than the other ones, indicating that the inflammatory storm was closely correlated with disease severity ^19–21^. The pro-inflammatory cytokines are known to not only perpetuate the inflammation to impair lung function, but also perturb bone metabolism, leading to bone resorption ^18,22^. A recent radiological study on COVID-19 survivors with persisting symptoms for up to three months after discharge revealed that the inflammation in bone marrow persisted after recovery ^23^. Additionally, reactive hemophagocytosis mediated by cytokine storm-induced activation of macrophage was common in deceased COVID-19 patients ^24^. Therefore, we hypothesize that in addition to other reported extrapulmonary manifestations, the cytokine storm in severe COVID-19 may also contribute to pathological changes in the skeletal system ^2^.

In this study, we characterized the previously unreported effects of SARS-CoV-2 infection on bone metabolism during the acute and post-recovery phases in our established golden Syrian hamster model which closely mimics human infection ^25^. Moreover, we identified inflammation-induced osteoclastic activation as the underlying mechanism mediating the pathological bone loss. The findings of this study highlight the need for optimizing clinical protocols for monitoring long-term complications of COVID-19 and finding novel treatment strategies for SARS-CoV-2-induced inflammatory osteopenia / osteoporosis.

## Results

### SARS-CoV-2 causes significant loss of bone trabeculae

To investigate the *in vivo* effects of SARS-CoV-2 infection on bone metabolism, we utilized our established golden Syrian hamster model, which recapitulates the clinical, virological, immunological, and pathological features of COVID-19 in humans ^25^ (**Fig. 1a & 1b**). In this hamster model, the most prominent disease manifestations are seen at about 4 days post-infection (dpi) and the hamsters generally recover at about 7 to 10 dpi. As shown in our three-dimensional micro computerized tomography (μCT) scans, SARS-CoV-2-infected but not PBS-challenged mock-infected control hamsters exhibited progressive loss of bone trabeculae at the distal metaphysis of femurs from the acute phase (4 dpi) to the post-recovery phase (30 dpi) and chronic phase (60 dpi) of infection (**Fig. 1c**). A significant decrease in the thickness of trabeculae was detected at as early as 4 dpi (**Fig. 1d**). Quantitatively, dramatic reductions (up to 50%) in trabecular bone volume fraction, trabecular number, and polar moment of inertia were seen at 30 dpi. There was also significant decrease in bone mineral density and trabecular thickness, as well as significant increase in the specific bone surface and trabecular pattern factor at 30 dpi. The bone volume and density then remained static from 30 dpi to 60 dpi. To verify our μCT scan findings, we examined the histological changes of the femurs of SARS-CoV-2-infected and mock-infected hamsters (**Fig. 1e, Extended Data Fig. 1a**), which demonstrated that the reduction in bone trabeculae structures at the distal metaphysis of femur was evident at both 30 dpi and 60 dpi in the SARS-CoV-2-infected but not mock-infected hamsters.

**Figure 1:**
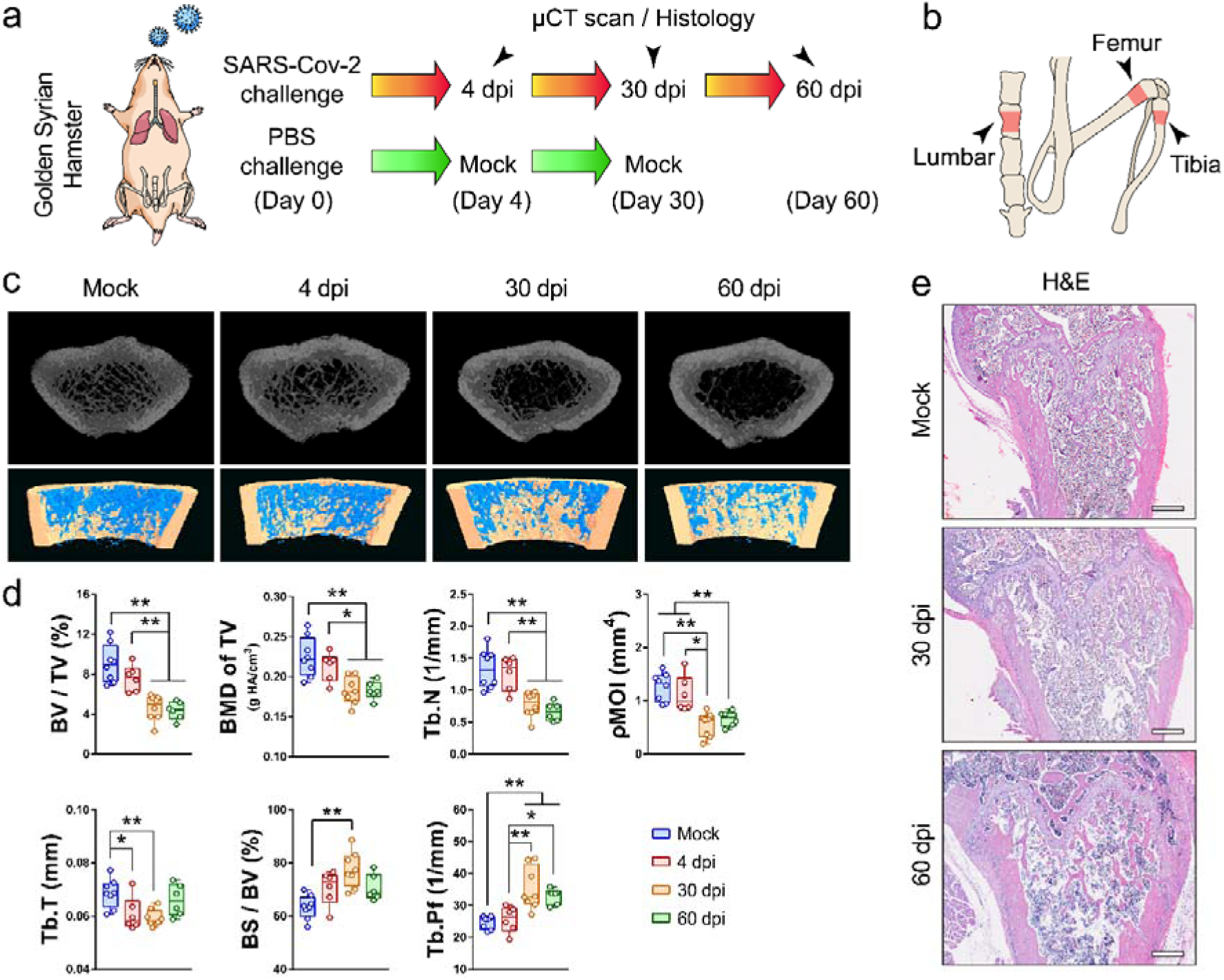
SARS-CoV-2 induces bone loss in the golden Syrian hamster model. **(a)** Golden Syrian hamsters were either challenged with SARS-CoV-2 or PBS (Mock), followed by μCT scan and histology analysis at 4, 30 and 60 days post-infection (dpi). **(b)** The regions of interest for μCT evaluation of bone density included the distal metaphysis of femur, proximal metaphysis of tibia, and lumbar vertebrae. **(c)** Representative μCT images showing the reduction in trabecular bone volume in the femurs of the SARS-CoV-2-infected hamsters. **(d)** Corresponding measurements of trabecular bone volume fraction (BV/TV), bone mineral density (BMD of TV), trabecular number (Tb.N), polar moment of inertia (pMOI), trabecular thickness (Tb.T), specific bone surface (BS/BV), and trabecular pattern factor (Tb.Pf). **(e)** Representative H&E staining images showing the cancellous bone structures in the femurs of the SARS-CoV-2-infected hamsters (scale bars = 500 µm). Data are mean□±□SEM. ns: *P*>0.05, \**P*<0.05, \*\**P*<0.01 by one-way ANOVA with Tukey’s *post hoc* test.

Next, we investigated whether similar changes were seen in other sites. Both the μCT scan data and histological analysis showed that there was also significant decrease in bone trabeculae at the proximal metaphysis of tibia (**Fig. 2a to 2c, Extended Data Fig. 1a**). The trabecular bone volume of SARS-CoV-2-infected hamsters at 30 dpi and 60 dpi was only about 50% of that of the mock-infected hamsters (**Fig. 2b**). Corroboratively, there was decreased bone mineral density and trabecular number, and increased trabecular pattern factor at 30 dpi and 60 dpi. In contrast, the changes in trabecular bone thickness, polar moment of inertia, and specific bone surface were evident at as early as 4 dpi. Similar to our findings in the femur, the bone structures remained relatively static between 30 dpi and 60 dpi. The same pattern of bone loss with significant reduction in trabecular bone volume fraction, bone mineral density, and trabecular thickness, and higher specific bone surface and trabecular pattern factor were observed also in the lumbar vertebrae at 30 dpi (**Fig. 2d & 2e**). Overall, these findings showed that SARS-CoV-2 infection causes significant bone loss at different sites of the skeleton in the hamster model.

**Figure 2:**
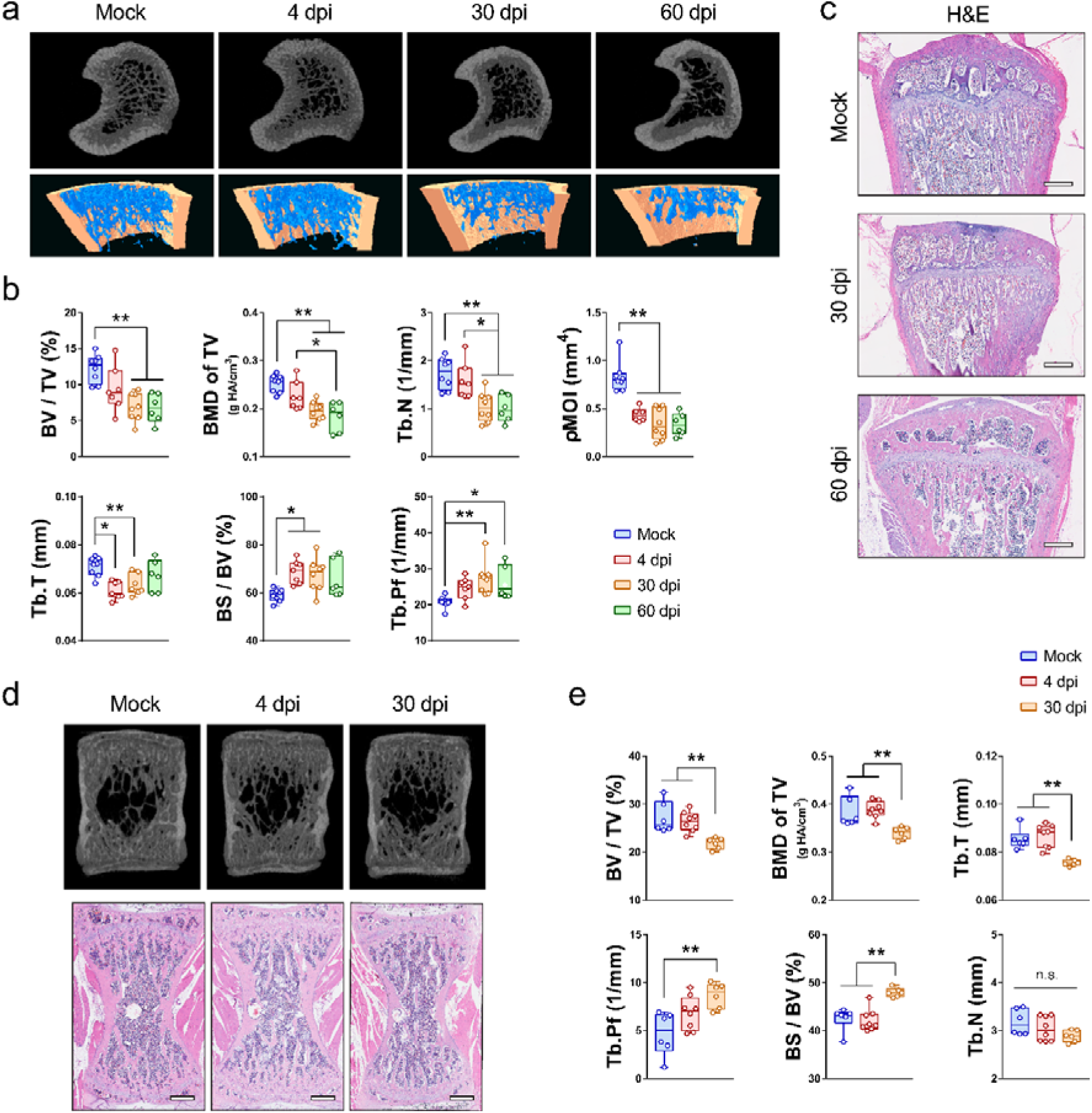
Bone loss in the tibias and lumbar vertebrae after SARS-CoV-2 infection in the golden Syrian hamster model. **(a)** Representative μCT images showing the reduction in trabecular bone volume in tibias of the SARS-CoV-2-infected hamsters. **(b)** Corresponding measurements of BV/TV, BMD of TV, Tb.N, pMOI, Tb.T, BS/BV, and Tb.Pf. **(c)** Representative H&E staining images showing the cancellous bone structures in tibias of the SARS-CoV-2-infected hamsters (scale bars = 500 µm). **(d)** Representative μCT images showing the bone loss in the lumbar vertebrae of the SARS-CoV-2-infected hamsters. **(e)** Corresponding measurements of BV/TV, BMD of TV, Tb.N, pMOI, Tb.T, BS/BV, and Tb.Pf. Data are mean□±□SEM. ns: *P*>0.05, \**P*<0.05, \*\**P*<0.01 by one-way ANOVA with Tukey’s *post hoc* test.

### SARS-CoV-2 activates osteoclastogenesis in hamsters

To provide mechanistic insights into the dysregulated bone metabolism in SARS-CoV-2-infected hamsters, we asked whether the bone loss is primarily caused by an alteration in bone resorption or bone formation. Compared with mock-infected hamsters, a significantly higher number of tartrate-resistant acid phosphatase-positive (TRAP^+^) osteoclasts were found in the bone trabeculae at the distal metaphysis of the femur (**Fig. 3a**), the proximal metaphysis of the tibia (**Fig. 3b**), and the lumbar vertebrae (**Fig. 3c**) of SARS-CoV-2-infected hamsters. The number of TRAP^+^ osteoclasts at the distal metaphysis of the femur of the SARS-CoV-2-infected hamsters at 4 dpi and 30 dpi was almost double of that of the mock-infected hamsters (**Fig. 3d**). Moreover, immunofluorescence staining demonstrated significantly more TRAP^+^ osteoclasts expressing nuclear factor of activated T-cells, cytoplasmic 1 (NFATc1) at the bone surface of SARS-CoV-2-infected hamsters (**Fig. 3e**). The increased intensity of TRAP and NFATc1 at the distal metaphysis of femur at 4 dpi indicated that the osteoclastic activity was upregulated by SARS-CoV-2 infection (**Fig. 3f**). We also compared the expression of NFATc1, TRAP, cathepsin K, and receptor activator of NF-κB (RANK) in the bone tissues using western blotting, showing these osteoclastic markers are upregulated after SARS-CoV-2 infection (**Extended Data Fig. 1b &1c)**. In contrast, ALP staining (**Extended Data Fig. 2a**) and immunofluorescence staining for osteocalcin (**Extended Data Fig. 2b**) showed that there was no significant difference in the osteoblastic activities between the SARS-CoV-2-infected and mock-infected hamsters at 4 dpi.

**Figure 3:**
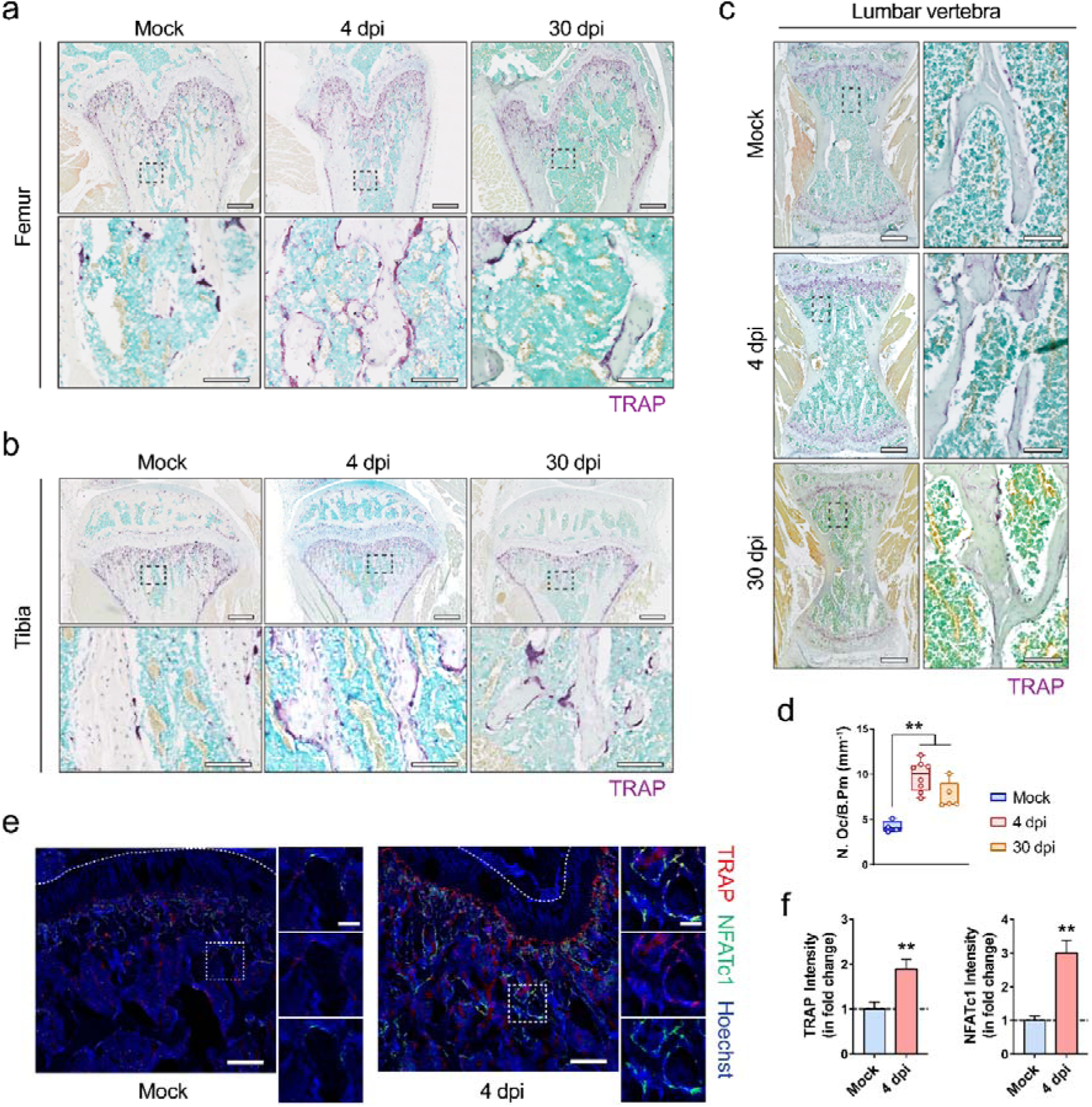
Osteoclastogenesis in the femurs and tibias of the hamsters after SARS-CoV-2 infection. **(a, b, c)** Representative TRAP staining showing the increase in the number of TRAP^+^ osteoclasts at **(a)** the distal metaphysis of femur, **(b)** the proximal metaphysis of tibia, and **(c)** the lumbar vertebrae after the challenge with PBS (Mock) or SARS-CoV-2 (4 dpi). Lower images (scale bars = 100 µm) are high-resolution versions of the boxed regions in the upper images (scale bars = 500 µm). **(d)** Corresponding quantification of TRAP^+^ osteoclasts at the trabecular bone surface of the golden Syrian hamsters four days after challenge with PBS (Mock) or SARS-CoV-2 (4 dpi). **(e)** Representative immunofluorescence staining images and **(f)** the corresponding fluorescence intensity quantification showing the increase in the number of TRAP^+^ NFATc1^+^ osteoclasts at the distal metaphysis of femur at 4 dpi. Tile scans (scale bars = 200 µm) of the distal femoral metaphysis are shown along with high-magnification of the boxed regions (scale bars = 50 µm). Data are mean□±□SEM. ns: *P*>0.05, \**P*<0.05, \*\**P*<0.01 by one-way ANOVA with Tukey’s *post hoc* test **(d)** or Student’s t-test **(f)**.

We next determined the expression of various osteoclastogenesis-related genes in the bone tissues of SARS-CoV-2-infected hamsters. Compared with mock-infected hamsters, the expression of receptor activator of nuclear factor-kappa B ligand (*RANKL*), which is essential for the osteoclastic differentiation, was tripled in SARS-CoV-2-infected hamsters (**Fig. 4a**). Moreover, several osteoclastic marker genes contributing to the formation and activity of osteoclasts, including receptor activator of NF-κB (*RANK*), cathepsin K (*CTSK*), matrix metallopeptidase 9 (*MMP-9*), and colony-stimulating factor 1 receptor (*CSF1R*), were significantly upregulated in the bone tissues after SARS-CoV-2 infection (**Fig. 4a**). The expression of osteoprotegerin (*OPG*), which acts as a decoy receptor for RANKL to inhibit RANK-RANKL mediated osteoclastogenesis and bone resorption, was correspondingly significantly downregulated after the infection (**Fig. 4a**). In addition to the increased number of TRAP^+^ osteoclasts, we further found significant higher number of osteoclast progenitors, including CD68^+^ macrophages and RANK^+^ preosteoclasts, in SARS-CoV-2-infected hamsters (**Fig. 4b & 4c**). There were more multinuclear cells located at the bone trabeculae co-expressing TRAP and CD68 in the SARS-CoV-2-infected hamsters (**Fig. 4b**). Using multiplex immunohistochemical (IHC) staining, we further identified this lineage of TRAP^+^RANK^+^CD68^+^ osteoclasts upregulated in response to SARS-CoV-2 infection to be expressing a higher level of interleukin-1 beta (IL-1β) than the ones seen in the mock-infected hamsters (**Fig. 4d**). Thus, to further elucidate the underlying mechanism, we then examined the expression of genes related to IL-1β signaling. Compared with the mock-infected hamsters, SARS-CoV-2-infected hamsters exhibited significant upregulation of interleukin-1 receptor type I (*IL-1RI*), but not interleukin-1 receptor type II (*IL-1RII*). Interestingly, the expression of IL-1β in the bone tissue was not significantly different between the two groups (**Fig. 4e**). Moreover, SARS-CoV-2 infection also induced a two-fold increase in the expression of interleukin-1 receptor antagonist (*IL-1RA*) in bone tissue. Taken together, these findings indicate SARS-CoV-2 infection leads to the activation of osteoclastic cascade resulting in the destruction of trabeculae structure in both long bones and axial skeleton.

**Figure 4:**
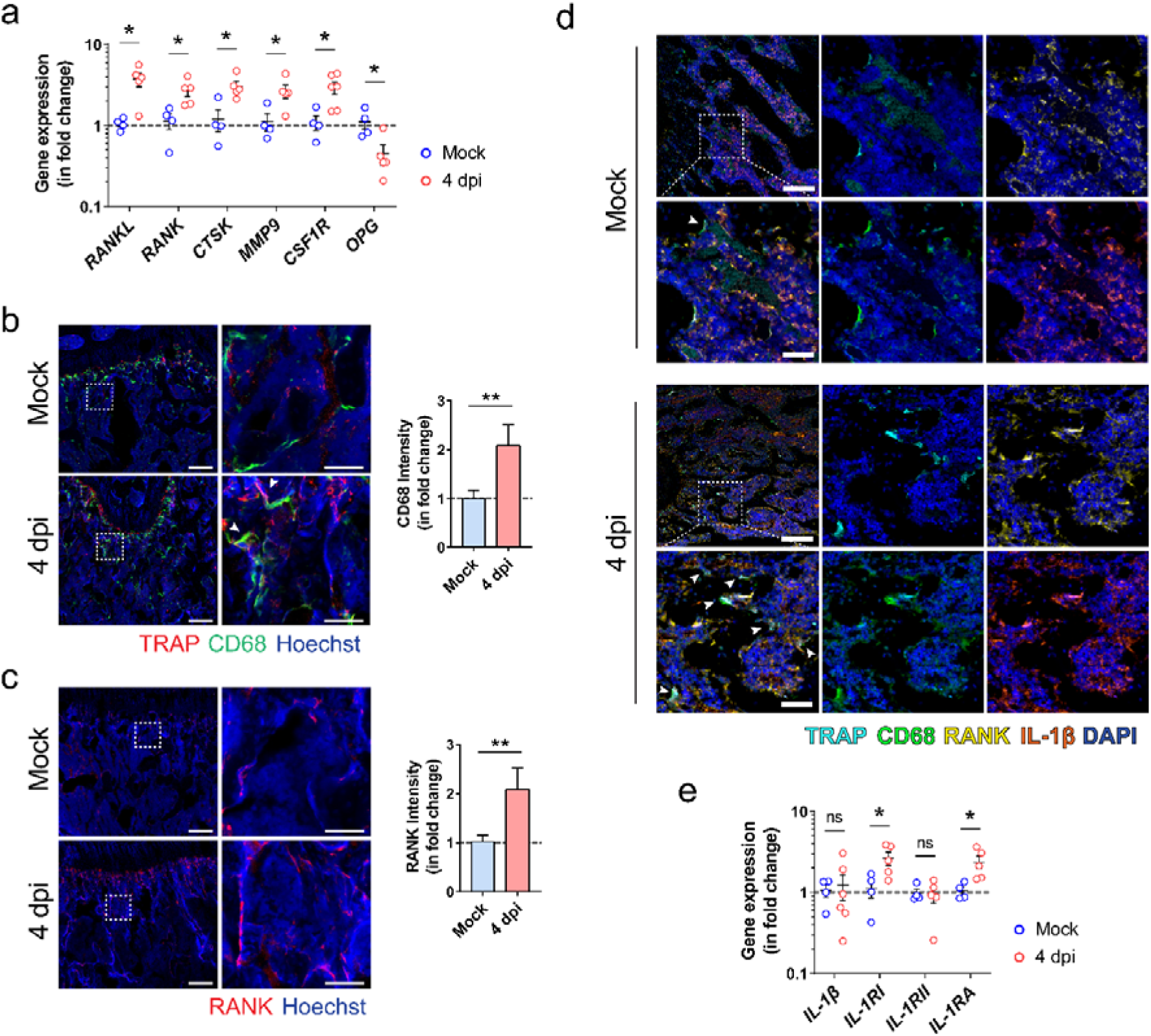
The inflammatory activation and osteoclastic differentiation of monocyte-macrophage lineage after SARS-CoV-2 infection. **(a)** The expression of osteoclastogenesis-related genes in the bone tissue of the hamsters at day 4 after challenge with either PBS (Mock) or SARS-CoV-2 (4 dpi). **(b)** Representative immunofluorescence staining images showing the increase in the number of CD68^+^ macrophages and TRAP^+^ osteoclasts after SARS-CoV-2 infection. Tile scans (scale bars = 200 µm) and high-magnification of the boxed regions (scale bars = 50 µm), as well as corresponding quantification for the fluorescence intensity of CD68 are shown. **(c)** Representative immunofluorescence staining images and the corresponding quantification showing the upregulation of RANK-expressing cells at the trabecular bone surface after SARS-CoV-2 infection. Tile scans (scale bars = 200 µm) and high-magnification of the boxed regions (scale bars = 50 µm) are shown. **(d)** Representative multi-colour immunohistochemical staining for TRAP, CD68, RANK, and IL-1β was performed at the distal metaphysis of femur at day 4 after challenge with either PBS (Mock) or SARS-CoV-2 (4 dpi). DAPI was used for nuclear counterstaining. Tile scans (scale bars = 200 µm) and high-magnification of the boxed regions (scale bars = 50 µm) are shown. **(e)** The expression of IL-1β signaling-related genes in bone tissue at day 4 after challenge with either PBS (Mock) or SARS-CoV-2 (4 dpi). Data are mean□±□SEM. ns: *P*>0.05, \**P*<0.05, \*\**P*<0.01 by Student’s t-test.

### SARS-CoV-2 disturbs inflammatory microenvironment in skeleton system without direct infection

Having demonstrated the involvement of the local immune response in pathological osteoclastogenesis in SARS-CoV-2 infection, we next asked whether the inflammatory bone loss was also caused by direct infection of the bone tissues by SARS-CoV-2. In SARS-CoV-2-infected hamsters, viral nucleocapsid protein (NP) in co-localized angiotensin-converting enzyme 2 (ACE2)- expressing cells were evident throughout the respiratory tract, from the nasal turbinate to the trachea and pulmonary alveoli at 4 dpi (**Extended Data Fig. 3**). CD68^+^ macrophages engulfing SARS-CoV-2-infected cells, which co-expressed viral NP and ACE2, were also observed, indicating active immune response in the areas. In stark contrast, despite the presence of ACE2 in some of the immune cells residing in bone tissues, viral NP was absent in the periosteum, bone trabeculae, and synovium of the femoral bone tissue in SARS-CoV-2-infected hamsters (**Fig. 5a**). Moreover, viral RNA was not detected in the bone tissue (**Fig. 5b**). These findings indicated that the bone tissues were not directly infected by SARS-CoV-2. We then investigated whether the osteoclastogenesis was associated with the virus-induced inflammatory response. Our ELISA results showed that the mean serum IL-1β, tumor necrosis factor-α (TNF-α), and IL-6 protein levels were all higher in the SARS-CoV-2-infected hamsters than that of the mock-infected hamsters, with the levels of IL-1β and TNF-α reaching statistical significance (P<0.01) (**Fig. 5c**).

**Figure 5:**
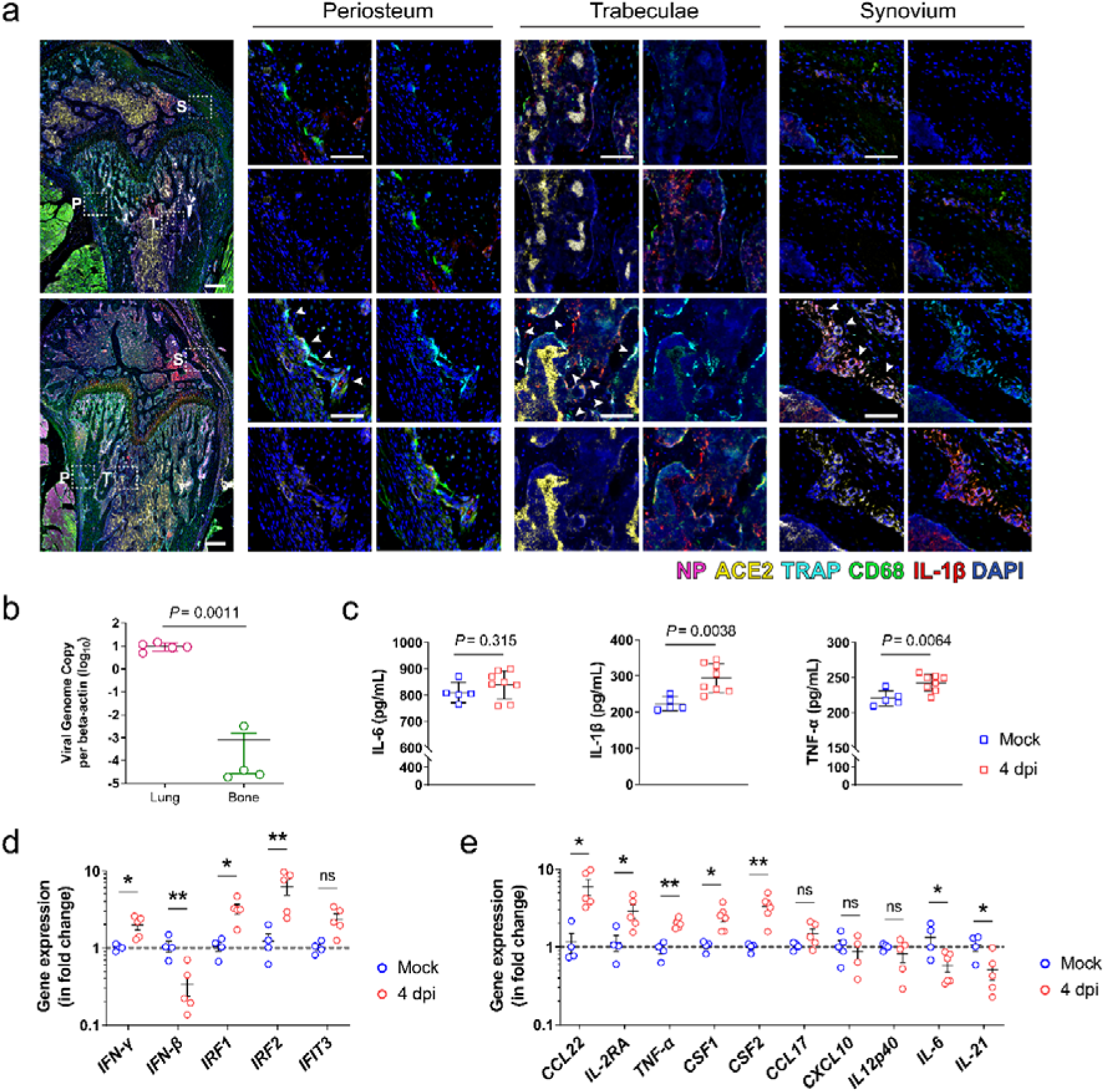
The absence of SARS-CoV-2 infection of bone tissues. **(a)** Representative multi-colour immunohistochemical staining for SARS-CoV-2 nucleocapsid protein (NP), angiotensin-converting enzyme 2 (ACE2), TRAP, CD68, and IL-1β was performed at the distal metaphysis of femur after challenge with either PBS (Mock) or SARS-CoV-2 (4 dpi). Tile scans (scale bars = 200 µm) and high-magnification of the boxed regions (P, periosteum; T, trabeculae; S, synovium; scale bars = 50 µm) are shown. **(b)** Viral genome copies of the lung tissue (n=5) and bone tissue (n=4) harvested from SARS-CoV-2 infected hamsters. **(c)** The inflammatory cytokines, including IL-6, IL-1β, and TNF-α, in the serum of the hamsters challenged with either PBS (Mock) or SARS-CoV-2 (4 dpi). **(d)** The expression of interferon signaling-related genes and **(e)** viral infection-associated inflammatory genes in bone tissue at day 4 after challenge with either PBS (Mock) or SARS-CoV-2 (4 dpi). Data are mean□±□SEM. ns: *P*>0.05, \**P*<0.05, \*\**P*<0.01 by Student’s t-test.

Interestingly, while direct SARS-CoV-2 infection of bone tissue was absent, the expression of interferon-gamma (*IFN-*γ) and its downstream signals, including interferon regulatory factor 1 (*IRF1*) and interferon regulatory factor 2 (IRF2), were significantly upregulated in the bone tissue of SARS-CoV-2-infected hamsters at 4 dpi (**Fig. 5d**). The expression of interferon-induced protein with tetratricopeptide repeats 3 (*IFIT3*) was only marginally increased. Compared with the mock control, the inflammation-related genes upregulated in bone tissue in response to the respiratory infection of SARS-CoV-2 include C-C motif chemokine 22 (*CCL22*), interleukin-2 receptor antagonist (*IL-2RA*), *TNF-*α, colony stimulating factor 1 (*CSF1*), and colony stimulating factor 2 (*CSF2*), nevertheless, the expression of C-C motif chemokine 17 (*CCL17*), C-X-C motif chemokine ligand 10 (*CXCL10*), and interleukin 12 p40 (*IL12p40*) remained similar (**Fig. 5d & 5e**). Additionally, it is noteworthy that the infection in the respiratory system intriguingly downregulated the expression of interferon-beta (*IFN-*β), interleukin 21 (*IL-21*), and interleukin 6 (*IL-6*) in bone tissue, which are vital for the clearance of virus infection.

### SARS-CoV-2-induced inflammatory cytokines upregulate osteoclastogenesis

To address whether the bone loss subsequent to SARS-CoV-2 infection was caused by the pro-inflammatory cytokines that originate from the respiratory system, we first compared the expression of various inflammatory cytokines in the bone tissue of SARS-CoV-2-infected and mock-infected hamsters. Using immunofluorescence staining, we showed that SARS-CoV-2 infection contributed to a significant increase in the levels of IL-1β, IL-1RA, and TNF-α, as well as a marginal increase in the level of IFN-γ in bone tissue (**Fig. 6a to 6d**). These changes were confirmed by Western blot which demonstrated a more than six-fold increase of IL-1β, a seven-fold increase of TNF-α, and a three-fold increase of IL-1RA in bone tissue at 4 dpi in the SARS-CoV-2-infected hamsters than the mock-infected hamsters (**Extended Data Fig. 4a & 4b**). Additionally, immunofluorescence staining showed the co-expression of these inflammatory cytokines with CD68, the bone resident macrophage marker (**Fig. 6a to 6d**). More importantly, we showed that the expression of NF-κB, the key transcription factor in inflammatory responses, was significantly higher at the bone surface of SARS-CoV-2-infected hamsters than that of the mock-infected hamsters (**Fig. 6e**). Semi-quantification by both immunofluorescence staining and Western blotting showed that the level of NF-κB in the bone tissue was doubled in the bone tissues of SARS-CoV-2-infected hamsters than the mock-infected hamsters (**Fig. 6a, Extended Data Fig. 4a & 4b**).

**Figure 6:**
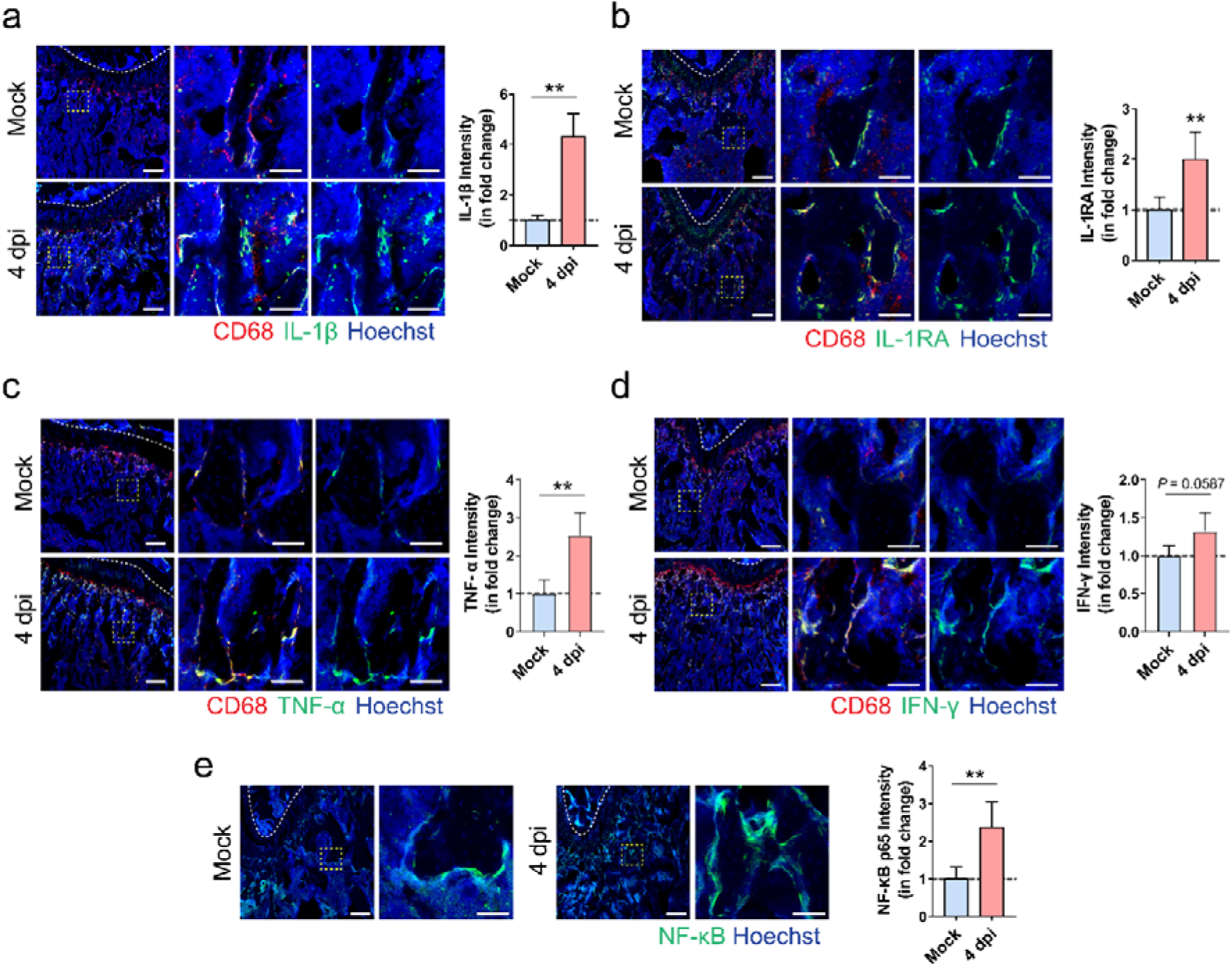
Macrophage-mediated inflammatory response in bone tissues after SARS-CoV-2 infection. **(a-d)** Representative immunofluorescence staining images and the corresponding quantification showing the increase in the expression of **(a)** IL-1β, **(b)** IL-1RA, **(c)** TNF-α, and **(d)** IFN-γ, at the distal metaphysis of femur after challenge with either PBS (Mock) or SARS-CoV-2 (4 dpi). Tile scans (scale bars = 200 µm) and high-magnification of the boxed regions (scale bars = 50 µm) are shown. **(e)** Representative immunofluorescence staining images and the corresponding quantification showing the increase in NF-κB p65 expression at the distal metaphysis of femur after challenge with either PBS (Mock) or SARS-CoV-2 (4 dpi). Data are mean□±□SEM. \*\**P*<0.01 by Student’s t-test.

To further confirm the effects of these inflammatory cytokines on osteoclastic activities in bone tissue, we conducted a series of experiments using mouse bone marrow macrophages (BMMs) isolated from young (three-month-old) or adult (six-month-old) mice. First, we tested the response of murine BMM to the stimulation of IL-1β as our data had shown the activation of the IL-1β signaling cascade in SARS-CoV-2-infected hamsters (**Fig. 4e)**. The presence of recombinant murine IL-1β contributed to a two-fold increase in the number of TRAP^+^ multinuclear cells derived from BMM isolated from young mice (**Fig. 7a & 7b**). When added to BMMs isolated from adult mice, IL-1β led to a doubled size of fused osteoclasts (**Fig. 7a & 7b**). In contrast, the addition of IL-1β neutralizing antibody (Neu-Abs) not only reduced the number of BMMs-derived osteoclasts, but also resulted in a smaller size of osteoclasts differentiated from adult mouse BMMs. Meanwhile, the expression of *IL-1R1* was increased by six times in the young BMMs and four times in the adult BMMs **(Fig. 7c)**. However, the effect of Neu-Abs on the downregulation of *IL-1R1* was only significant in BMMs from young mice, but not the ones from adult mice. Nevertheless, the gene expression of *IL-1RA*, which encodes interleukin-1 receptor antagonist, was significantly upregulated in both young and adult BMMs in response to the stimulation of IL-1β and downregulated after the addition of Neu-Abs.

**Figure 7:**
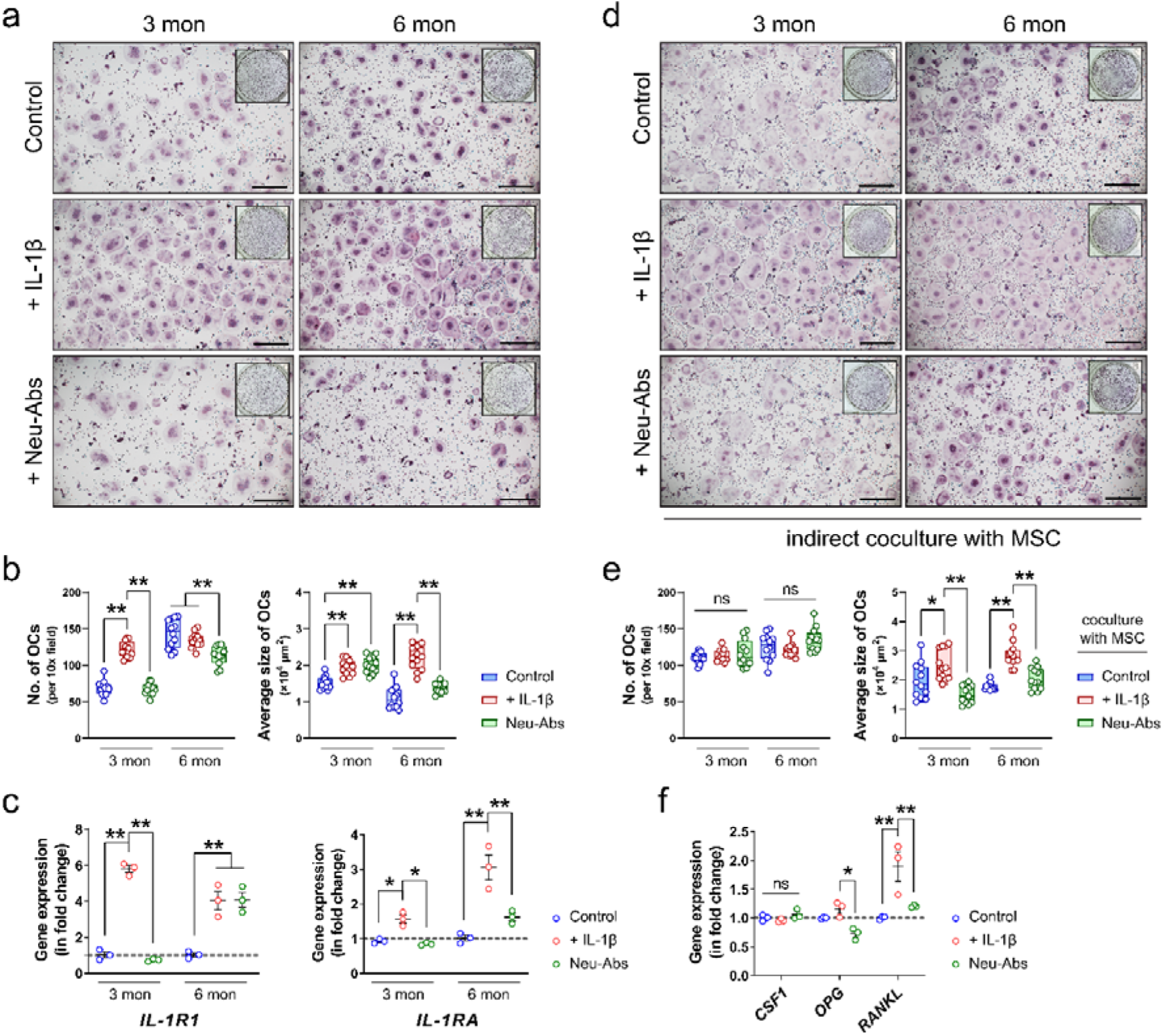
Inflammatory cytokine IL-1β promotes osteoclastogenesis. **(a)** Representative microscopic images (scale bars = 500 µm) and **(b)** the corresponding quantifications showing the formation of TRAP^+^ multinuclear cells from bone marrow macrophage (BMM) isolated from three-month-old young mice (3 mon) and six-month-old adult mice (6 mon). Recombinant murine IL-1β or its neutralizing antibody (Neu-Abs) was added to the culture medium throughout the osteoclastic induction using M-CSF and RANKL. **(c)** The gene expression of *IL-1R1* and *IL-1RA* in BMMs from young or adult mice with or without the presence of recombinant murine IL-1β or its neutralizing antibody. **(d)** Representative microscopic images (scale bars = 500 µm) and **(e)** the correspond quantifications showing the formation of TRAP^+^ multinuclear cells when the BMMs were indirectly co-cultured with mesenchymal stem cells (MSCs) stimulated with recombinant murine IL-1β or its neutralizing antibody. **(f)** The gene expression of *CSF1, OPG*, and *RANKL* in MSC-treated with recombinant murine IL-1β or its neutralizing antibody. Data are mean□±□SEM. ns: *P*>0.05, \**P*<0.05, \*\**P*<0.01 by two-way ANOVA with Tukey’s *post hoc* test.

Besides the direct effect of IL-1β on BMMs, we further tested whether IL-1β could promote osteoclastogenesis through the regulation of the osteoblast lineage. The indirect co-culture of BMMs and MSC was achieved using a transwell assay. IL-1β led to significant increase in the size of BMM-derived osteoclasts after their co-culture with IL-1β-treated MSC, even though the number of TRAP^+^ multinuclear cells remained unchanged (**Fig. 7d & 7e**). Neu-Abs-treated MSC, instead, decreased the average size of osteoclast derived from young or adult BMMs. This might be explained by the changes in the pro-osteoclastogenesis cytokines produced by MSCs, as IL-1β contributed to a two-fold increase in the expression of *RANKL* without changing the level of *OPG* in MSC (**Fig. 7f**). Neither IL-1β nor its neutralizing antibody significantly regulated the expression of *CSF1* in MSC.

The effects of IL-1β and its neutralizing antibody on osteoclastogenesis were further confirmed using Western blotting. In both osteoclasts derived from BMM of young mice and that of adult mice, the expressions of NFATc1, NF-κB p65, and CTSK were significantly upregulated by recombinant murine IL-1β (**Fig. 8a**). Additionally, Neu-Abs suppressed the pro-osteoclastogenesis effects of IL-1β, as it downregulated the expression of NFATc1, NF-κB p65, and CTSK in BMM-derived osteoclasts from young and adult mice. Moreover, the Neu-Abs inhibited the IL-1β induced phosphorylation of c-Jun N-terminal kinase (JNK) (**Extended Data Fig. 4c**). Similar to the direct effect of IL-1β on osteoclastogenesis, IL-1β-treated MSC also led to a significant upregulation in the expression of NFATc1 and the phosphorylation of JNK, which were both significantly downregulated by Neu-Abs-treated MSC (**Extended Data Fig. 5a**). Although IL-1β-treated MSC only contributed to a marginal increase in the expression of NF-κB p65 and CTSK in BMM, Neu-Abs-treated MSC significantly inhibited both of them (**Extended Data Fig. 5a**).

**Figure 8:**
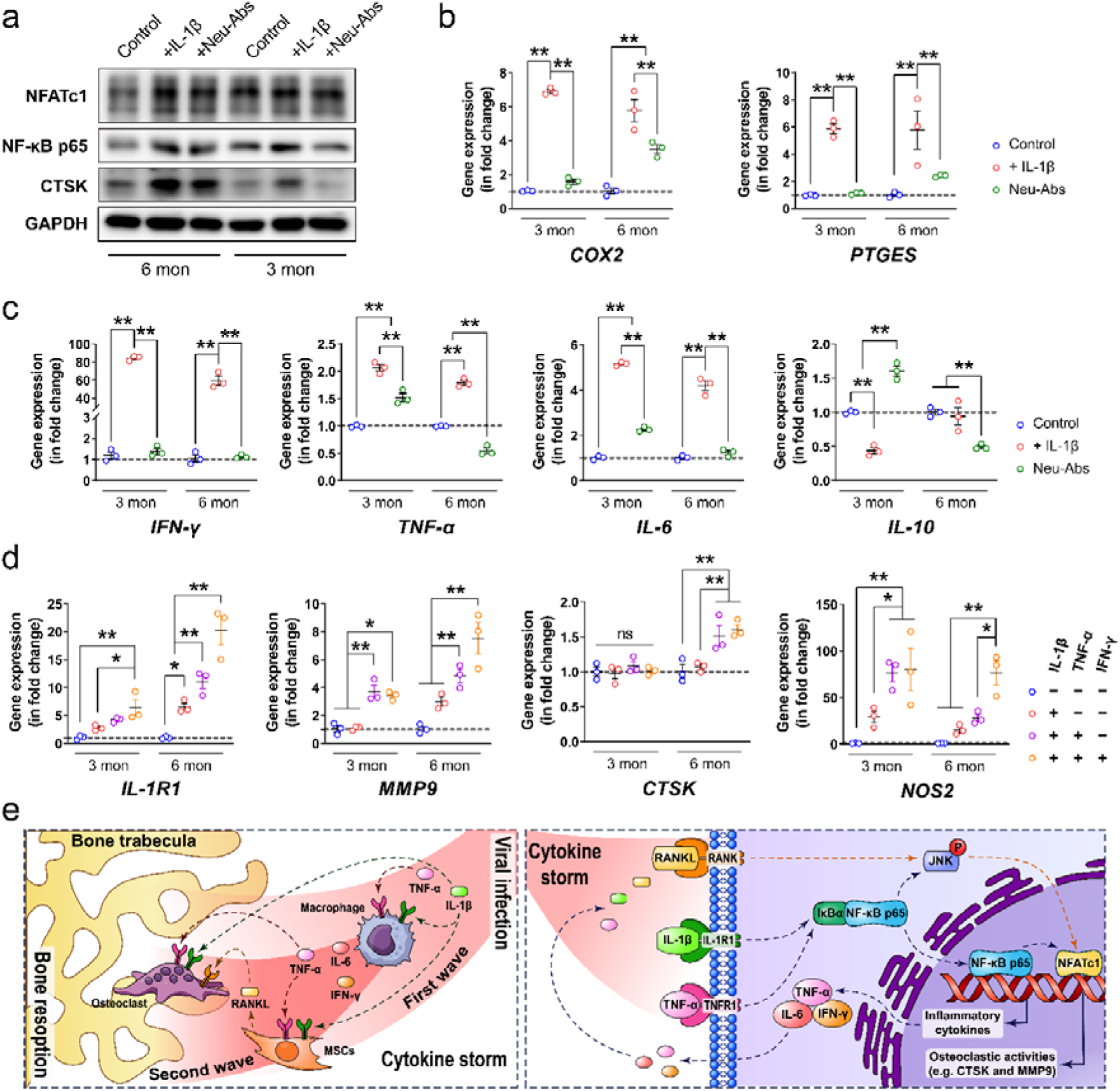
Inflammatory cytokines exacerbates the pro-inflammatory response in bone marrow macrophages. **(a)** Representative Western blots showing the expression of NFATc1, NF-κB p65, and CTSK in bone marrow macrophages (BMMs) isolated from adult (6-mon) or young (3-mon) mice with or without the presence of recombinant murine IL-1β or its neutralizing antibody. **(b)** The gene expression of pain associated cytokines and **(c)** other pro/anti-inflammatory cytokines in BMMs from young or adult mice with or without the addition of recombinant murine IL-1β or its neutralizing antibody. **(d)** The effects of various inflammatory cytokines on the gene expression of *IL-1R1, MMP9, CTSK*, and *NOS2*. **(e)** Schematic diagram showing the mechanism through which the systemic cytokine storm induced by viral infection of the respiratory tract exacerbates the local inflammatory response in bone tissue, leading to pathological bone resorption. Data are mean□±□SEM. ns: *P*>0.05, \**P*<0.05, \*\**P*<0.01 by two-way ANOVA with Tukey’s *post hoc* test.

### The second wave of cytokine storm promotes pathological bone resorption

Since our results have verified the absence of SARS-CoV-2 infection in bone tissue, we hypothesized the prominent inflammatory response in bone tissue was related to the systemic cytokine storm that originates from the respiratory system after viral infection. Therefore, in addition to the pro-osteoclastogenesis effect of IL-1β, we also explored the immunomodulatory effects of IL-1β and its neutralizing antibody on BMM isolated from young and adult mice. The recombinant murine IL-1β contributed to an approximately 6-fold increase in the expression of *COX2* and *PTGES* (**Fig. 8b**), which are both involved in the synthesis of prostaglandin E2 (PGE2). The presence of Neu-Abs inhibited the increase in the expression of *COX2* and *PTGES* induced by IL-1β. The pro-inflammatory effect of IL-1β can also be manifested by the significant changes in the expression of various inflammation-related genes. For instance, in BMM isolated from young mice, IL-1β led to a more than 80-fold increase in *IFN-*γ expression and an around 2-fold increase in *TNF-*α expression, as well as a 5-fold increase in *IL-6* expression. In contrast, the expressions of *IL-10* and *IL-23* were significantly downregulated by IL-1β, with the expression of IL-22 remained unchanged (**Fig. 8c, Extended Data 5b**). In BMM isolated from adult mice, IL-1β also contributed to a 60-fold increase in *IFN-*γ expression, an approximately 1.75-fold increase in *TNF-*α expression, an around 4-fold increase in *IL-6* expression, and a less than 1.5-fold increase in *IL-22* expression. However, the expression of *IL-10* and *IL-23* were not significantly altered by the addition of IL-1β (**Fig. 8c, Extended Data 5b**). In BMM from both young and adult mice, the addition of Neu-Abs abolished the upregulation in the expression of *IFN-*γ, *TNF-*α, and *IL-6* caused by IL-1β. In BMM from young mice, Neu-Abs resulted in significant upregulation in the expression of *IL-10, IL-22*, and *IL-23* (**Fig. 8c, Extended Data 5b**).

We then asked whether the pro-inflammatory cytokines from the systemic cytokine storm subsequent to SARS-CoV-2 infection or the activated macrophages in bone tissue synergistically promoted osteoclastogenesis. First, we showed that recombinant murine TNF-α and IFN-γ exaggerated the effects of IL-1β through the upregulation of *IL-1R1* expression (**Fig. 8d**). This phenomenon was more prominent in BMM isolated from adult mice, as IL-1β, TNF-α, and IFN-γ together led to an around 20-fold increase in the expression of *IL-1R1*, while IL-1β alone only upregulated *IL-1R1* by 5-fold. As a result, two major marker genes (i.e., *MMP9* and *CTSK*) for osteoclastic activities were found to be further upregulated when the three inflammatory cytokines were administered in combination (**Fig. 8d**). In BMM isolated from young mice, the addition of TNF-α and IFN-γ to IL-1β resulted in a 4-fold increase in *MMP9* expression, when IL-1β alone failed to induce higher *MMP9* expression than the control. Meanwhile, in BMM isolated from adult mice, the combination of IL-1β, TNF-α, and IFN-γ led to a more than 7-fold increase in the expression of *MMP9* and a 1.5-fold increase in the expression of *CTSK*, while IL-1β alone did not significantly alter the expression of these two osteoclastic marker genes. Notably, the three inflammatory cytokines synergistically contributed to a more than 60-fold increase in the expression of Nitric Oxide Synthase 2 (*NOS2*), when IL-1β alone only upregulated *NOS2* expression by less than 30 times. Together, these findings indicate that the pro-inflammatory cytokines synergistically contribute to pathological bone resorption.

## Discussion

In addition to respiratory tract manifestations, extrapulmonary manifestations are also commonly reported in severe coronavirus infections such as COVID-19, SARS, and MERS ^4,26,27^. Based on the phylogenetic similarities of SARS-CoV-2 and SARS-CoV, it has been postulated that the two betacoronaviruses may cause similar clinical features in infected patients. However, recent evidence has increasingly shown that there are more musculoskeletal sequelae associated with COVID-19 than SARS ^10,28^. The most severe musculoskeletal complication in SARS patients was non-progressive avascular necrosis of the femoral head caused by high-dose steroid pulse therapy ^29^. In contrast, musculoskeletal sequelae have been increasingly reported in severe COVID-19 patients including those who have recovered from the acute phase of the infection ^10,28^. In this study, using an established hamster model, we demonstrated significant bone resorption at the acute inflammatory stage after SARS-CoV-2 infection. Moreover, in addition to a significantly increased number of RANK^+^ osteoclast precursors at this stage ^30^, there were also more TRAP^+^ osteoclasts expressing NFATc1, which is known to serve as a master regulator for terminal differentiation of osteoclasts ^31^. This implies that pathological bone destruction may happen quickly after the onset of COVID-19. More importantly, similar findings were evident in different bone tissues harvested from the hamsters, suggesting that the bone loss is not site-specific but systemic. Without proper intervention, the bone volume and bone mineral density were barely restored even after the viral load became undetectable at the post-recovery / chronic inflammatory phase. The presence of pathological bone loss may in turn complicate the rehabilitation of COVID-19 patients. For example, low bone mineral density is a known risk factor for vertebral fractures that may impair the respiratory function of COVID-19 patients in the rehabilitation phase ^32,33^. It was recently reported that thoracic vertebral fractures occurred in 36% of COVID-19 patients and increased the patients’ need for noninvasive mechanical ventilation ^34^.

Osteoporosis, which is characterized by decrease in bone mass, microarchitectural bone disruption, and skeletal fragility leading to higher fracture rate, has been extensively reported in critically ill patients ^35^. The inflammatory cytokines are suggested to be one of the most vital mediators of the pathological bone loss that occurs in these diseases because they not only activate osteoclasts, but also impede osteoblast function ^18^. Cytokine dysregulation has been associated with various clinical manifestations of COVID-19, including some involving the musculoskeletal system, such as myalgia, sarcopenia, arthralgia, and arthritis ^36,37^. Mechanistically, our findings in this study demonstrated that SARS-CoV-2 induced pathological bone resorption through two waves of cytokine storms instead of direct infection in the skeletal tissue. Since we didn’t detect significant upregulation in gene expression of *IL-1*β from bone tissue, the increase in IL-1β level in bone tissue may primarily attribute to the SARS-CoV-2-induced immune response in the respiratory system. After the first wave of inflammatory cytokines (e.g., IL-1β and TNF-α) reaches skeletal tissue via circulation, they quickly modulated the monocyte-macrophage lineage residing there to initiate a second wave of cytokine storm in the bone tissue. The second wave of cytokines produced in the skeletal system, including but not limited to IFN-γ, IL-6, and PGE2, not only contribute to osteoclastogenesis in several interdependent signaling pathways ^18^, but also augment the pro-inflammatory actions of IL-1β and TNF-α by upregulating the expression of IL-1R1, which was implicated to primarily mediate pathological bone resorption ^38^.

Among the various inflammatory factors that may be associated with osteoclastogenesis ^22^, we identified IL-1β and TNF-α as the key mediators for SARS-CoV-2-induced bone loss in the hamster model. As two of the most significantly upregulated inflammatory cytokines in the serum samples of SARS-CoV-2-infected hamsters, IL-1β and TNF-α synergistically contribute to the pro-inflammatory microenvironment in the skeletal tissue, leading to inflammatory bone loss. IL-1β and TNF-α have been respectively reported to play essential roles in RANKL-induced osteoclast formation ^39,40^. Besides, they are also demonstrated to be interdependent in mediating inflammatory osteopenia ^41^. We found that the concurrence of three pro-inflammatory cytokines (i.e., IL-1β, TNF-α and IFN-γ) in the bone tissue after SARS-CoV-2 infection preferably upregulated *IL-1R1*, which is primarily expressed in pathologically activated osteoclasts responsible for inflammatory bone destruction ^42,43^. In contrast, the expression of *IL-1R2*, which serves as a decoy receptor for IL-1β to negatively regulate IL-1β signalling, was not significantly altered. Consequently, these SARS-CoV-2-induced pro-inflammatory cytokines dramatically upregulate the expression of *MMP9* and *CTSK*, which are both known to play dominant roles in the degradation of extracellular matrix ^44^. Additionally, we also found that the inflammatory cytokines elevated in COVID-19 (e.g., IL□1β) were able to promote the formation of osteoclasts via regulating the production of RANKL from MSC, as similarly reported in other conditions ^45^. Importantly, we demonstrated that BMMs isolated from young mice were more responsive to the pro-osteoclastic stimulation of IL□1β. This is clinically relevant because pediatric patients generally have stronger ability to adequately respond to viral infections with rapid production of high levels of pro-inflammatory cytokines ^46,47^. This heightened pro-inflammatory response, together with the lower baseline pro-inflammatory state in children, makes them more susceptible to various syndromes related to immune dysregulation ^47,48^. However, we also find that the involvement of other pro-inflammatory cytokines, such as TNF-α and IFN-γ, contributed to a more prominent effect on promoting the osteoclastic activities in BMMs isolated from adult mice than the ones from young mice. This might explain why musculoskeletal symptoms are mostly seen in adult patients other than children and the elderly ^49^.

Besides the immunomodulatory effect and the pro-osteoclastogenesis effect, the accumulation of various pro-inflammatory cytokines in the skeletal tissue can lead to several other long-term health concerns. For example, we showed that IL□1β dramatically upregulated *COX2* and *PTGES*, which both contribute to the production of PGE2 ^50^. PGE2 is not only an inflammatory mediator involved in bone modeling but also a neuromodulator that can sensitize peripheral sensory neurons leading to inflammatory pain ^51^. Therefore, the lasting (from week 0 to week 28) bone ache or burning feeling reported by more than 20% of COVID-19 patients ^10^ indicates long-term monitoring of the inflammatory status of bone tissue after the recovery of the disease might be necessary. The inflammation in bone tissue can also alter the output of immune cells or cytokines from the bone marrow, which are supposed to participate in combating viral infection ^52^. In our study, the significantly upregulated expression of *IFN-*γ and its signaling-associated genes (e.g., *IRF1* and *IRF2*) in bone tissue in response to the cytokine storm indicate they might play a role in protecting bone tissue from virus infection ^53^. However, it is also noteworthy that the expression of several vital anti-viral chemokines and cytokines, such as IFN-β ^54^, IL-21 ^55^, CCL17 ^56^, remained unchanged or even downregulated in the bone tissue. Thus, the suppression in the bone marrow-derived anti-viral factors may be one of the immune evading mechanisms for SARS-CoV-2 that warrants further investigation.

## Conclusions

In this study, we demonstrated the influence of SARS-CoV-2 infection on systemic bone loss during the acute and post-recovery / chronic phases. We revealed the cytokine storm derived from the respiratory system as the major contributor to pathological bone resorption. IL-1β and other pro-inflammatory cytokines disrupt the balance in bone metabolism and trigger a second wave of cytokine storm in the skeletal tissue to further augment their pro-osteoclastogenesis effect (**Fig. 8e**). The findings in our study highlight the need to closely monitor COVID-19 patients’ bone density. The benefits of prophylactic interventions against the development of pathological bone loss in COVID-19 patients should be further evaluated in animal models and clinical trials.

## Methods

### Virus and Biosafety

SARS-CoV-2 (strain HKU-001a, GenBank accession number: MT230904) was a clinical strain isolated from the nasopharyngeal aspirate specimen of a COVID-19 patient in Hong Kong ^57^. The plaque purified viral isolate was amplified by one additional passage in VeroE6 cells to make working stocks of the virus as described previously ^58^. All experiments involving live SARS-CoV-2 followed the approved standard operating procedures of The University of Hong Kong (HKU) Biosafety Level-3 facility ^59,60^.

### Animal Model

The animal experiments were approved by the HKU Committee on the Use of Live Animals in Teaching and Research. Briefly, 6–10-week-old male golden Syrian hamsters (*Mesocricetus auratus*) were obtained from the Chinese University of Hong Kong Laboratory Animal Service Centre through the HKU Centre for Comparative Medicine Research. The animals were kept in Biosafety Level-2 housing and given access to standard pellet feed and water ad libitum until virus challenge in our Biosafety Level-3 animal facility. Each animal was intranasally challenged with 10^5^ PFU of SARS-CoV-2 in 100 µl of PBS under intraperitoneal ketamine (200 mg/kg) and xylazine (10 mg/kg) anesthesia at 0 dpi as we previously described ^61^. Mock-infected animals were challenged with 50 µL of PBS. Their blood, bone and lung tissues were collected at sacrifice at 4, 30, and/or 60 dpi for µCT, virological, and histopathological analyses.

### Micro-CT analysis

The PBS or virus-challenged hamsters were sacrificed at 4, 30, and 60 dpi for micro-CT analysis of bilateral femurs and tibias. Before being transferred from the biosafety Level-3 facility, the specimens were fixed in 4% paraformaldehyde for 48 h and 70% ethanol for 24 h to inactivate the pathogens. The femurs and tibias were scanned by a high-resolution micro-CT scanner (SkyScan 1076, Kontich, Belgium) at a resolution of 11.53 μm per pixel. The voltage of the scanning procedure was 88 kv with a 110-μA current. Two phantom contained rods with standard density of 0.25 and 0.75 g/cm3 were used for calibration of bone mineral density (BMD). Data reconstruction was done using the NRecon software (Skyscan Company), the image analysis was done using CTAn software (Skyscan Company), and the 3D model visualization was done using CTvox (Skyscan Company) and CTvol (Skyscan Company). The bone volume fraction (BV/TV), specific bone surface (BS/BV), bone mineral density (BMD of TV), trabecular thickness (Tb. T), trabecular number (Tb. N), trabecular pattern factor (Tb.Pf) were measured by the µCT data.

### Histological analysis

Histology and immunohistochemical staining were performed on both paraffin sections and cryosections. In brief, the bone specimens, after fixation in 4% PFA for 48 hours, were decalcified with 12.5% ethylenediaminetetraacetic acid (EDTA, Sigma-Aldrich) for 4 weeks. For paraffin sections, the specimens were processed, embedded in paraffin, and cut into 5 µm-thick sections using a rotary microtome (RM215, Leica Microsystems, Germany). Haematoxylin and eosin (H&E) staining, TRAP staining (Sigma-Aldrich), and ALP staining (Sigma-Aldrich) were performed on selected sections from each sample following manufacturer’s instructions. Images were captured using the Vectra Polaris Imaging System (Akoya Biosciences, USA). For immunostaining, the samples were dehydrated in 20% sucrose solution with 2% polyvinylpyrrolidone (PVP, Sigma-Aldrich) for 24h and embedded in 8% gelatin (Sigma-Aldrich) supplemented with 20% sucrose and 2% PVP. Forty-μm-thick coronal-oriented sections of the femurs were obtained using a cryostat microtome. Immunostaining was performed using a standard protocol. Briefly, after blocking with 10% goat serum, the sections were incubated with primary antibodies to CD68 (Abcam, ab31630/ab125212), RANK (Abcam, ab13918), TRAP (Abcam, ab216025), osteocalcin (TAKARA, M186), IL-1β (Abcam, ab9722), IL-1RA (Abcam, ab124962), TNF-α (Abcam, ab9635), IFN-γ (Abcam, ab9657), NF-κB (CST, #8242), anti-NFATc1 (CST, #8032) overnight at 4°C. Alexa-Fluor 488-conjugated and Alexa-Fluor 647-conjugated secondary antibodies (Thermo Fisher Scientific) were used for immunofluorescent staining, while the nuclei were counterstained with Hoechst 33342 (Thermo Fisher Scientific). Immunofluorescent images were captured using LSM 780 confocal microscopy (Zeiss, Germany).

### Multiplex IHC analysis

Antigen retrieval and blocking were performed on the dewaxed slides using the Antigen retrieval reagent (pH 6.0) and Blocking/antibody diluent provided in the Opal Polaris Multicolor Manual IHC Detection Kit (Akoya Biosciences, USA), following the manufacturer’s instructions. In brief, the incubation of each primary antibody was done overnight at 4°C. The primary antibodies used in this study included anti-CD68 (Abcam, ab31630, USA), anti-TRAP (Abcam, ab216025), antii-RANK (Abcam, ab13918), anti-IL-1β (Abcam, ab9722), anti-NP (ThermoFisher, USA), anti-ACE2 (ThermoFisher). Between each incubation of the primary antibody, tyramide signal amplification (TSA) visualization was performed using the Opal Polymer Horseradish peroxidase (HRP) secondary antibody and fluorophores: Opal 520, Opal 570, Opal 620, Opal 690, and DAPI (Akoya Biosciences, USA). The stained slides were imaged using the Vectra Polaris Imaging System (Akoya Biosciences, USA).

### Cell culture

The mesenchymal stem cells (MSCs) and bone marrow macrophages (BMMs) were isolated from the long bones of 3-month-old or 6-month-old C57L6/J mice. In brief, the mice were euthanized with overdosage of intraperitoneal injection of Pentobarbital. After the removal of attached soft tissues using forceps and gauze, the dislocated femurs and tibias were dissected into pieces. The whole bone marrow cells were resuspended in a serum-free DMEM medium (Gibco, USA) using a vortex mixer, while the bone chips and debridement were removed by passing the mixture through a cell strainer. After centrifugation, the cell pellet was resuspended in a DMEM medium, supplemented with 10% FBS and 1% Penicillin-Streptomycin (complete DMEM medium), and cultured in culture flasks. After a 6-h incubation in a humidified incubator with 5% CO_2_ at 37°C, the unattached cells were gently removed for the induction of BMMs. Meanwhile, the attached bone marrow cells were gently washed in PBS and further cultured as MSCs in a complete DMEM medium until they reached 80% confluence. The culture medium was refreshed every 2 days, and only passages 3–5 were used for the experiments. After the 3-day macrophage induction using a complete DMEM medium supplemented with 20 ng/ml of macrophage colony-stimulating factor (M-CSF), the BMMs became adherent for the osteoclastic differentiation using a complete DMEM supplemented with 50 ng/ml of receptor activator of nuclear factor kappa-B ligand (RANKL, R&D System, USA) and 20 ng/ml of M-CSF (R&D System, USA). The coculture of BMMs with MSCs was done using a transwell system (0.4 µm pore size, Corning Costar, USA). In brief, MSCs were seeded in the transwell inserts and cultured overnight with 5% CO_2_ at 37°C for attachment. Afterwards, the transwell inserts were placed into the lower chambers in which BMMs have been differentiated using medium supplemented with M-CSF for three days as described previously. To mimic the challenge of pro-inflammatory microenvironment in COVID-19, 1 ng/mL recombinant murine IL-1b (R&D system, USA) was supplemented in the medium. For the inhibition of IL-1β, 10 ng/mL IL-1b neutralizing antibody (Abcam, ab9722) was added to the cell culture. For studying the synergistic effects of pro-inflammatory cytokines on BMMs, the culture medium of BMMs was further supplemented with 1 ng/mL TNF-α (R&D system, USA) and 1 ng/mL IFN-γ (R&D system, USA).

### Real-time quantitative polymerase chain reaction (RT-qPCR) assay

The total RNA from the bone specimens were isolated using Trizol reagent (Thermo Fisher, USA) following the manufacturer’s instructions. The total RNA from the cultured cells was extracted and purified using an RNeasy Plus kit (Qiagen, USA) following the manufacturer’s instructions. For the reverse transcript, complementary DNA was synthesized using a Takara RT Master Mix (Takara, Japan), following the manufacturer’s instructions. The primers used in the RT-qPCR assay were synthesized using Integrated DNA Technologies (IDT, Singapore), based on sequences designed using Primer-BLAST (National Center for Biotechnology Information, NCBI, Table S1) or retrieved from the Primer Bank (http://pga.mgh.harvard.edu/primerbank/, Table S2). The SYBR Green Premix Ex Taq (Takara, Japan) was used for the amplification and detection of cDNA targets on the LightCycler480 Real-time PCR system (Roche, USA). The mean cycle threshold (Ct) value of each target gene was normalized to the housekeeping genes (i.e., RPL18 or GAPDH). The results were shown in a fold change using the ΔΔCt method.

### Western blotting

The proteins from homogenized bone tissue from the hamsters were isolated using Trizol reagent (Thermo Fisher, USA) following the manufacturer’s instructions. The proteins from cultured BMMs were harvested using RIPA Lysis and Extraction Buffer (ThermoFisher, USA) containing 1% Phosphatase Inhibitor Cocktail (ThermoFisher, USA). The concentration of protein was measured using the BCA Protein Assay Kit (ThermoFisher, USA). A total of 20 µg of protein from each sample was subjected to sodium dodecyl sulfate polyacrylamide gel electrophoresis (SDS-PAGE) and transferred to the polyvinylidene difluoride (PVDF) membrane (Merck Millipore, USA). Then, the membrane was blocked in 5% w/v bovine serum albumin (BSA, Sigma-Aldrich, USA) and incubated with blocking buffer-diluted primary antibodies overnight at 4°C. The primary antibodies used included mouse anti-NFATc1 (Santa Cruz, USA), mouse anti-TRAP (Abcam), rabbit anti-Cathepsin K (Abcam), mouse anti-RANK (Abcam), rabbit anti-NF-κB p65 (CST), rabbit anti-IL-1β (Abcam), rabbit anti-IL-1RA (Abcam), rabbit anti-TNF-α (Abcam), rabbit anti-phospho-JNK (CST), rabbit anti-JNK (CST), rabbit anti-β-actin (Abcam), The protein bands were visualized by using HRP conjugated secondary antibodies and an enhanced chemiluminescence (ECL) substrate (Advansta, USA) and exposed under the Typhoon5 Biomolecular Imager 680 (GE Amersham, USA).

### ELISA assay

The serum samples of the hamsters were collected at 4 dpi for cytokine/chemokine analysis. The serum level of IL-1β, TNF-α and IL-6 were detected using specific ELISA kit (MyBiosource, USA) following the manufacturer’s instructions.

### Statistical analysis

All data analyses were performed and illustrated using the Prism software (version 7, GraphPad, USA). The results were expressed as means ± standard error of the mean (SEM). The exact sample size (n) for each experimental group was clearly shown as dot plots in the figures and indicated in the legends. For comparisons among multiple groups, a one-way or a two-way analysis of variance (ANOVA) was used, followed by Tukey’s multiple-comparison *post hoc* test. The levels of significant difference among the groups were defined and noted as \**P* < 0.05 and \*\**P* < 0.01. The sample size was decided based on preliminary data, as well as on observed effect sizes.

## Acknowledgments

We sincerely thank the staff members at the Faculty Core Facility, Li Ka Shing Faculty of Medicine, The University of Hong Kong, the Centre for Comparative Medicine Research of The University of Hong Kong, and the Laboratory Animal Service Centre of The Chinese University of Hong Kong for their facilitation of the study. This work is partly supported by funding from the National Key R&D Program of China (R&D#2018YFA0703100, 2020YFA0707500 and 2020YFA0707504); the General Research Fund of Hong Kong Research Grant Council (Nos. 17207719, 17214516, and N_HKU725/16); Health and Medical Research Fund (No.19180712 and COVID1903010-7), the Food and Health Bureau, The Government of the Hong Kong Special Administrative Region, China; the Hong Kong Innovation Technology Fund (ITS/287/17 and ITS/405/18); Shenzhen Science and Technology Funding (JSGG20180507183242702); HKU-SZH Fund for Shenzhen Key Medical Discipline (SZXK2020084); Sanming Project of Medicine in Shenzhen, China (SZSM201612055 and SZSM201911014); the Science and Technology Commission of Shanghai Municipality (No. 18410760600); the International Partnership Program of Chinese Academy of Sciences (GJHZ1850); the Guangdong Financial Fund for High-Caliber Hospital Construction (174-2018-XMZC-0001-03-2125/D-10); the High Level-Hospital Program, Health Commission of Guangdong Province, China; the Innovation and Technology Fund (ITF), the Government of the Hong Kong Special Administrative Region; the Consultancy Service for Enhancing Laboratory Surveillance of Emerging Infectious Diseases and Research Capability on Antimicrobial Resistance for Department of Health of the Hong Kong Special Administrative Region Government; the Major Science and Technology Program of Hainan Province (ZDKJ202003); the research project of Hainan academician innovation platform (YSPTZX202004); and the Hainan talent development project (SRC200003). We are also grateful for the donations of May Tam Mak Mei Yin, Richard Yu and Carol Yu, the Shaw Foundation of Hong Kong, Michael Seak-Kan Tong, Lee Wan Keung Charity Foundation Limited, Hui Ming, Hui Hoy and Chow Sin Lan Charity Fund Limited, Chan Yin Chuen Memorial Charitable Foundation, Tse Kam Ming Laurence, Marina Man-Wai Lee, the Hong Kong Hainan Commercial Association South China Microbiology Research Fund, the Jessie & George Ho Charitable Foundation, Perfect Shape Medical Limited, Kai Chong Tong, Foo Oi Foundation Limited, Betty Hing-Chu Lee, Ping Cham So, and Lo Ying Shek Chi Wai Foundation. The funding sources had no role in the study design, data collection, analysis, interpretation, or writing of the report.

## Author contributions

W. Qiao performed the *in vitro* and *in vivo* tests, as well as analyzed and interpreted the data. H. Lau and H. Xie contributed to the *in vitro* tests and helped with the data analysis. V.K.-M. Poon and C.C.-S. Chan contributed to the infection of hamsters and the collection of specimens. H. Chu, S. Yuan, T.T.-T. Yuen, K.K.-H. Chik, J.O.-L. Tsang, C.C.-Y. Chan, J.-P. Cai, and C. Luo performed experiments and analyzed data. K.-Y. Yuen and K.M.-C. Cheung provided funding and interpreted data. J.F.-W. Chan and K.W.-K. Yeung contributed to study design, data interpretation and supervision of the project. W. Qiao, J.F.-W. Chan and K.W.-K. Yeung wrote the manuscript, with input from all authors.

## Competing interests

The authors declare no competing interests.

## Notes

### Competing Interest Statement

The authors have declared no competing interest.

